# Social tolerance mediates social learning in wild red-fronted lemurs (Eulemur rufifrons) and ring-tailed lemurs (*Lemur catta*)

**DOI:** 10.1101/2025.09.18.677073

**Authors:** Sandro Sehner, Fanomezana Ratsoavina, Claudia Fichtel

## Abstract

Social learning is widespread in the animal kingdom, allowing individuals to acquire information from conspecifics. Because such learning often follows the social network within a population, species with different social structures and tolerance levels may show distinct learning trajectories. We tested this in four groups of wild, egalitarian red-fronted lemurs (*Eulemur rufifrons*, N = 34) and three groups of wild, hierarchically organized ring-tailed lemurs (*Lemur catta*, N = 35) using a social diffusion experiment with feeding boxes. We also conducted a co-feeding test of social tolerance. The feeding boxes could be opened by either lifting or pushing the box. In experimental conditions, we trained a demonstrator in one of the two techniques; in control conditions, no demonstrator was trained. More red-fronted lemurs learned to open the box than ring-tailed lemurs, and they learned faster. Both species preferred the lifting technique, but when pushing was the seeded technique, most red-fronted lemurs adopted it, whereas ring-tailed lemurs did not. Order of acquisition diffusion analysis showed that in both species, social learning followed the social network and was more likely than individual learning. Social tolerance predicted the proportion of learners within a group, with red-fronted lemurs being more tolerant than ring-tailed lemurs. In a similar vein, in the more tolerant red-fronted lemurs, scrounging occurred much more often than in the less tolerant ring-tailed lemurs. Scrounging might also have driven the preference for lifting technique in red-fronted lemurs but not in ring-tailed lemurs. Our results show that tolerance facilitates social learning and that information spreads along social network pathways. Most importantly, our results show that these effects vary across species with different social structures. Together, this suggests that social tolerance can drive group-specific behaviours and may play a key role in the evolution of cultural traits.

**Highlights:** **In the more socially tolerant red-fronted lemurs more individuals learned faster how to open a food box than in the less tolerant ring-tailed lemurs**

**The transmission of the opening technique followed the affiliative social network**

**Scrounging behaviour occurred more often in the socially tolerant red-fronted lemurs**

## Introduction

Social learning plays an essential role in the spread and maintenance of cultural traits (Boyd & Richerson, 1988; Tomasello et al., 1993; Whiten, 2021). Hence, understanding the mechanisms of effective social learning is crucial for our understanding of the roots of human cultural evolution. Social learning is not restricted to cohesive social groups, but it is more likely to occur in groups where individuals maintain close proximity (Coussi-Korbel & Fragaszy, 1995; Galef & Laland, 2005; van Schaik et al., 1999). It has been demonstrated that animals that are closely affiliated with each other are more likely to learn from each other (Garcia-Nisa et al., 2023) and that populations that exhibit more time in close association show a more variable cultural repertoire (Whiten & van Schaik, 2007). Thus, proximity-based observational learning requires a minimum level of social tolerance, broadly defined as the willingness to spend time in proximity with conspecifics and do so non-agonistically (De Waal, 1986; DeTroy et al., 2022; Rina Evasoa et al., 2019). For example, social tolerance, assessed at the beginning of a social diffusion experiment, predicted pathways of social learning in bearded capuchins (*Sapajus libidinosus*; Coelho et al. 2024). Though it remains unclear whether animals’ greater affiliation leads them to learn more effectively from each other, or whether learning from each other increases their affiliation (Karakoc et al., 2025; Kulahci et al., 2018).

One way to examine whether social learning is facilitated by proximities between individuals is via a ‘Network-based diffusion analysis’ (NBDA). This approach uses observations of social interactions and proximity to identify social networks, which can subsequently be analysed to examine how social learning occurs within these networks (Franz & Nunn, 2009). Since their initial introduction, NBDAs have been developed and extended (Hasenjager et al., 2021) and used across taxa to demonstrate that the diffusion of a socially learned trait is transmitted along the social network of a group via close associates (reviewed in Whiten et al., 2016). For example, in cetaceans, the spread of a foraging innovation, like ‘lobtail feeding’ in humpback whales (*Megaptera novaeangliae*), and sponge tool use in bottlenose dolphins (*Tursiops aduncus*), followed the structure of the social network (Allen et al., 2013; Wild et al., 2019). Equally, in primates, chimpanzees (*Pan troglodytes*), the spread of an innovation of using moss as a tool to soak water from a pond could also be tracked down along the social network (Hobaiter et al., 2014). Similar patterns have been found in vervet – and capuchin monkeys (*Chlorocebus pygerythrus*; *Cebus capucinus*), where social learning of various innovations regarding foraging followed the social network (Barrett et al., 2017; Canteloup et al., 2020, 2021). Moreover, a measure of social networks, Eigenvector centrality, which reflects the strength of the individual’s connection within a group, predicted the speed and probability of an individual acquiring a foraging technique in squirrel monkeys (*Saimiri sciureus*) (Claidière et al., 2013). Thus, patterns of close associates within groups reflect the spread of socially learned traits.

Accordingly, patterns of social transmission should also match variations in social structures between species. In other words, the degree of social learning is expected to correspond to a species level of social tolerance, with more egalitarian species exhibiting greater reliance on social learning and a higher degree of behavioural homogeneity compared to more despotic species, which may rely more on individual learning (Sehner et al., 2022). For instance, whereas humans heavily rely on social learning and evolved mechanisms like teaching for high-fidelity information transmission (O’Madagain & Tomasello, 2022; van Schaik et al., 2019; Whiten et al., 2022) and even imitate irrelevant actions when demonstrated by an adult, chimpanzees are more likely to develop individual solutions and only selectively learn socially (Gergely et al., 2002; Horner & Whiten, 2005). However, social learning is not restricted to species with high social tolerance, which can be seen across macaques. Macaques can be sorted alongside a tolerance spectrum (Thierry, 2007) and even less tolerant species like Japanese macaques (*Macaca fuscata*) and rhesus macaques (*M. mulatta*) have demonstrated social learning in various contexts (Garcia-Nisa et al., 2023; Kawai, 1965; Macellini et al., 2012). However, comparative studies examining differences in social learning are still rare.

So far, only a few studies have examined how the spread of socially learned traits varies between species exhibiting different social structures. For example, a comparative study between gorillas (*Gorilla gorilla gorilla*) and chimpanzees on learning a tool-use task revealed that fewer gorillas learned the task, whereas most chimpanzees acquired the task, with chimpanzees being the more tolerant species (Lonsdorf et al., 2009). Equally, more socially tolerant orangutans and chimpanzees rely on social learning to overcome food neophobia, whereas less socially tolerant gorillas do not (Gustafsson et al., 2014). In addition, more tolerant tufted capuchins (*Sapajus apella*) were more likely to find a solution to a series of extractive foraging devices, compared to less tolerant white-faced capuchins (Sehner et al., 2025). In contrast, a comparative study on social learning in baboons (*Papio ursinus*) and vervet monkeys (*Chlorocebus aethiops*) revealed higher levels of social learning in the less tolerant baboons, most likely because they displayed more intense social monitoring (Cambefort, 1981).

Interestingly, a comparative analysis of social networks among 78 primate groups (24 species) supports the notion that information transmission is affected by the structure of the social networks. Network optimality, i.e., how efficiently information is spread with a minimum number of social connections, is linked to cognitive abilities and social systems (Pasquaretta et al., 2014). In particular, network centralization and modularity affect efficiency, suggesting that social information in more egalitarian social systems with small differences in centrality and modularity between group members is more efficiently spread than in hierarchical systems (Pasquaretta et al., 2014). Yet, more empirical studies on the effect of the social network structure and social network metrics on social learning in species exhibiting different social systems are needed to shed light on the basic requirements for the development of traditions and ultimately on the evolution of culture.

In this study, we conducted a social diffusion experiment (Whiten & Mesoudi, 2008) to compare the spread of a socially learned trait in two highly comparable lemur species that differ in their social structure and levels of social tolerance among group members (Kappeler et al., 2022). To this end, we tested wild groups of rather egalitarian red-fronted lemurs (*Eulemur rufifrons*; Ostner & Kappeler, 2004; Pereira et al., 1990) and of hierarchically organized ring-tailed lemurs (*Lemur catta*; Pereira & Kappeler, 1997) in Madagascar. In red-fronted lemurs, a central male dominates other males, but subordinates cannot be further ranked in a linear hierarchy (Ostner & Kappeler, 1999). Moreover, contrarily to most other lemurs (Kappeler et al. 2022, females neither exhibit female dominance (Pereira et al., 1990) nor establish clear dominance relationships (Ostner & Kappeler, 1999). In ring-tailed lemurs, females are dominant over males, and both sexes exhibit clear dominance relationships (Pereira & Kappeler, 1997). Furthermore, both species have comparable body size, brain size, and diet (Isler et al., 2008; Johnson, 2007; Wilson & Hanlon, 2010), which makes them excellent models to address this question. We also know from previous studies that both species are capable of using social learning in field settings (red-fronted lemurs: Schnoell & Fichtel, 2012; but see Huebner & Fichtel, 2015; ring-tailed lemurs: Kendal et al., 2010) and that red-fronted lemurs exhibit higher social tolerance levels than ring-tailed lemurs (Fichtel et al., 2018). We predicted i) a larger number of individuals acquiring the task, ii) faster learning, and iii) more behavioural homogeneity within the groups of the more tolerant red-fronted lemurs in comparison to the less tolerant ring-tailed lemurs. In addition, social network metrics such as global efficiency, centralization, and modularity should affect the spread of information in the two species.

## Methods

### Study site and subjects

This study was conducted on two wild species of lemurs, red-fronted lemurs (*Eulemur rufifirons*) at the research station of the German Primate Center in Kirindy Forest, Western Madagascar, and ring-tailed lemurs (*Lemur catta*) in Berenty Private Reserve, Southern Madagascar. The experiments on four groups of wild red-fronted lemurs (groups A, B, F, and J) were conducted in 2011. All groups participated in other experimental studies before (Pyritz et al., 2013; Schnoell et al., 2014; Schnoell & Fichtel, 2012). Three groups of wild ring-tailed lemurs were tested in 2011 (group C1) and 2012 (groups C2A and YF). Group C2A was naïve to feeding experiments, but groups C1 and YF took part in an experimental study before (Kendal et al., 2010). All study subjects were well habituated to human presence due to long-term studies at the two sites (Kirindy Forest: Kappeler & Fichtel, Berenty Private Reserve: 2012; Jolly, 1966; Koyama et al., 2005) and could be individually identified either by nylon collars (red-fronted lemurs: Kappeler & Fichtel, 2012) or by their unique facial coloration (ring-tailed lemurs). Group size varied between 4 and 11 individuals in red-fronted lemurs and between 11 and 12 individuals in ring-tailed lemurs. A total of 34 red-fronted lemurs (14 females) and 35 ring-tailed lemurs (17 females) participated in the experiments.

### Feeding boxes and experimental procedure

Study subjects were confronted with food boxes (Figure 1a) that could be opened by using one of two techniques (push or lift; Figure 1b,c) to get access to the hidden reward (raisins).

**Figure 1:**
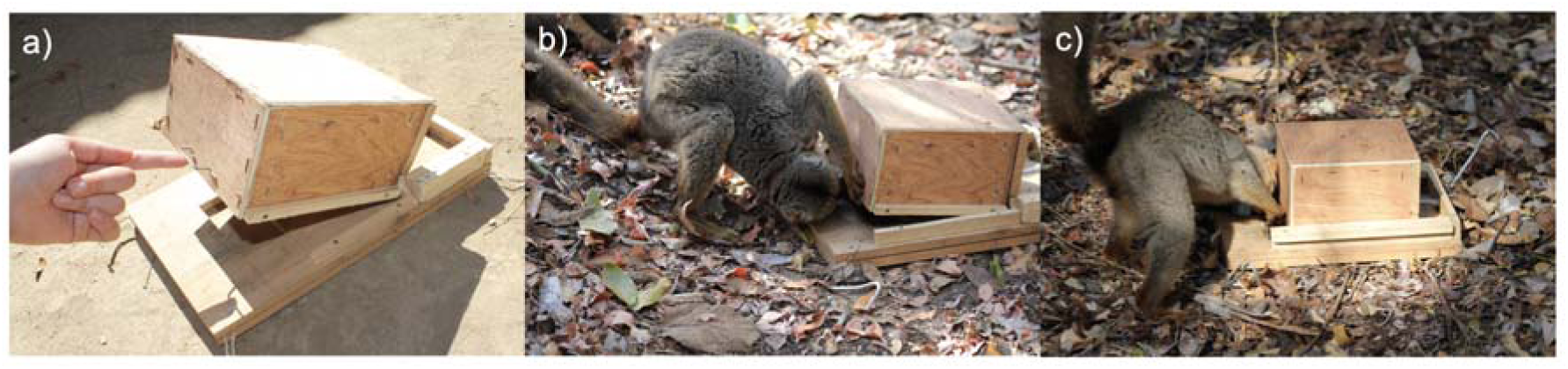
a) Food box that can be opened either by b) lifting or c) pushing.

To successfully open the food box, animals needed to make use of both hands, one hand to keep the box open and the other hand to reach for the reward. The food box consisted of a wooden board with a size of 30 cm x 20 cm x 2 cm on which a wooden box (size: 15 cm x 15 cm x 10 cm) was mounted. The board had a hollow carved inside, in which the reward was placed (Figure 1a). The wooden box had a handle attached at the front to allow better manipulation of the box (Figure 1a). When animals opened the feeding box with the lifting technique (Figure 1b), the box shut automatically afterward due to the weight of the upper part. When animals used the pushing technique, the box shut automatically afterward because a spring was installed at the back of the box that pushed the box back into its place (Figure 1d).

Before we conducted the experiment, we trained in two groups of red-fronted and ring-tailed lemurs a demonstrator for the lifting and pushing technique. Because individuals rarely separate from their social group, it was impossible to train one individual out of sight of the other group members. Therefore, we trained the demonstrator inside the social group by offering one constrained feeding box that would allow only one opening technique (see also van de Waal et al., 2010). Since in both species, usually one individual monopolized the food box, the rest of the group received a consistent demonstration of one of the two solutions. In red-fronted lemurs, an adult male (AMTho) in group A served as a demonstrator for the lifting technique and an adult male (FMCas) in group F as a demonstrator for the pushing technique. In ring-tailed lemurs, the dominant female (Viv) of group C1 served as a demonstrator for the lifting technique, and an adult female (Mil), the daughter of the dominant female of group YF, served as a demonstrator for the pushing technique. The training phase lasted until the demonstrator conducted at least 10 successful actions in four sessions each. A successful action consisted of opening the box and feeding on the reward. The demonstrators needed 6 to 7 sessions to reach this criterion, and the rest of the group thereby received at least 40 demonstrations of the opening technique before the experiment started. After the training phase of the demonstrator, we presented in the following 10 sessions three unconstrained boxes, where individuals could apply both techniques to open the box. Because a few individuals monopolized the three boxes, we presented five boxes during the subsequent 10 sessions.

Additionally, two groups in red-fronted lemurs (groups B and J) and one group in ring-tailed lemurs (group C1) served as control groups, in which we did not train a demonstrator and presented from the beginning on unconstrained boxes where the individuals could apply both techniques to open the boxes. As in the demonstrator groups, we presented three boxes in the first 10 sessions and five boxes in the subsequent 10 sessions. Experiments were video-taped with two cameras (Sony HDR-XR350 and a Sony DCR-SR75E camcorder). Feeding boxes were always baited before the groups approached the boxes to avoid that animals associate human observers with food. The experimental set-up was installed in some distance to the group, and the animals were attracted to the test location by using an auditory signal (Schnoell & Fichtel, 2012).

### Video analyses

Based on the video recordings, we recorded the time and technique of each action on the box, the identity of the individual manipulating the box, and whether the opener was able to retrieve the reward.

### Behavioural observations

We conducted in total 413 hours of animal focal observations in red-fronted lemurs (13 females and 23 males) and 579 hours (14 females and 13 males) in ring-tailed lemurs. Continuous focal sampling for 30 min and scan sampling every 10 min were used to collect behavioural data on the lemurs (Altmann, 1974). We employed a standard protocol (modified after Pereira & Kappeler, 1997) and used affiliative behaviours, i.e., time spent grooming and resting in body contact, to construct social networks and to calculate the social network metrics.

### Statistic analyses

For each species, we used a proportion test (function “prop.test”) to examine whether the proportion of individuals that learned the task differs from the proportion of individuals that did not learn the task. To assess factors influencing the probability that an individual learned the task, we used a binomial GLMM with a logit link with *learning* (yes/no) as a response. *Species*, the *presence of a demonstrator* (yes/no), and the interaction between the two factors, as well as *sex*, were used as fixed factors, and *age class* as a control factor. *Identity of the group* was included as a random factor. Because we offered at the beginning only three food boxes per group and group size varied more strongly between 4 and 11 individuals in red-fronted lemurs than in ring-tailed lemurs (11-12 individuals), resulting in differential access to the food boxes in groups with different *group sizes*, we included the log-transformed group size as an offset term. The model was fitted using the R package lme4 (Bates et al., 2014). The full model was compared to the null model, including only the control and random factors by using a likelihood ratio test (using the Anova function with argument test set to “χ²”).

The speed of learning was examined by comparing the number of trials until the first successful trial between species using a Mann-Whitney U-test. The influence of a demonstrator on learning speed was examined by comparing the number of trials until the first successful trial between groups with a demonstrator and open groups separately for each species using a Mann-Whitney U-test.

In order to test whether individuals were more likely to use the technique seeded by the demonstrator we fitted a binomial GLM with the proportion of using the lifting technique (using the “cbind” function (N trials using lifting technique, N trials using pushing technique) as response and treatments (demonstrator: lifting, demonstrator: pushing, no demonstrator) as a fixed factor for each species respectively. To examine whether one of the techniques might have been more difficult, we compared the proportion of unsuccessful actions when using the lifting or pushing technique with a Wilcoxon signed paired rank test for each species, respectively. Since we were not always able to reliably identify the used opening technique in 2.99% of openings (85 out of 2839 openings) in red-fronted lemurs and 0.91% of openings (47 out of 5143 openings) in ring-tailed lemurs, we removed these events from the dataset for these analyses.

The difference between species in proportion of scrounging events in relation to successful events was examined with a proportion test. To examine which factors influence the proportion of scrounging events, we fitted for each species a GLMM with logit-link function including the *proportion of successful scrounging events* using the “cbind” function with the number of successful scrounging events and the number of successful openings without scrounging. As predictors, we included *technique used* by, *age class* and *sex* of the solving individual as well as the z-transformed *session number*. We included *individual* and *group identity* as random factors and *session number* as random slopes with individual and group identity. We excluded the correlation among random slopes and intercepts of session number within group identity due to convergence issues.

To examine whether social tolerance and social network metrics influenced the proportion of learners within a group in both species, we ran a Spearman’s rank correlation. Definition of social network metrics: social tolerance: the average number of individuals feeding together on a clumped food resource (DeTroy et al., 2022; Fichtel et al., 2018). Global and average efficiency indices were taken from Pasquaretta et al. (2014) for all groups except red-fronted lemur group B, because it consisted only of four individuals and the sample size is too small to calculate the efficiency parameters. Efficiency is defined as how fast information can spread through a network with a minimum number of connections. It ranges from 0 – 1 with more efficient networks having values closer to 1. The first efficiency coefficient, global efficiency, is the ratio between the number of individuals divided by the number of connections multiplied by the network diameter (the longest of the shortest paths). The second coefficient, average efficiency, is computed by inversing the shortest path length for each pair of individuals within the network.

### Network-Based Diffusion Analysis (NBDA)

We also conducted a ‘Network-Based Diffusion Analysis’, which uses an association matrix, presenting the proportion of time that individuals are associated, assuming that the spread of a behaviour follows the social network structure (Franz & Nunn, 2009; Hoppitt et al., 2010). NBDA models examine the spread of a behaviour as a stochastic process in which, at any given time, each individual has a learning rate that determines the likelihood of learning the behaviour at that time (Hasenjager et al., 2021). We employed the Order of Acquisition Diffusion Analysis (OADA; Hoppitt et al., 2010), which is based on the assumption that the order of acquisition follows a social network. The different models were fitted to the data by maximum likelihood and tested against models with no social transmission by using the corrected Akaike’s Information Criterion for small sample size (AICc). Because some individuals were able to monopolize the food boxes, we included rates of aggression given and sex as potentially confounding factors to investigate their effects on learning rates. Time spent grooming and resting in body contact were used to construct the social network. We estimated in total six models. In model 1, social transmission was set constant. In model 2, we added the individual levels variables (ILVs), i.e., aggression rates and sex. In model 3, we allowed the magnitude of social transmission to vary between groups. In model 4, we added the ILVs to model 3. In model 5, we allowed the magnitude of social transmission to be constant within conditions (demonstrator and open groups) but to vary between conditions. In all models, we accounted for the fact that in some groups the diffusion was seeded by using the argument ‘demons’ indicating which individual served as a demonstrator. We used the R script model for NBDA Version 1.2.11 available at https://lalandlab.wp.st-andrews.ac.uk/freeware/.

Although a Time of Acquisition Diffusion Analysis (*TADA*) is usually statistically more powerful (Hasenjager et al., 2021; Hoppitt et al., 2010), we did not employ it for the following reasons. The *TADA* assumes that the rate of acquisition is at a given time is only dependent on the status of other individuals in the groups (Franz & Nunn, 2009; Hasenjager et al., 2021; Hoppitt et al., 2010). In our study, the first individuals who acquired the task monopolized the feeding boxes during the first 10 sessions, in which we presented only three boxes. After adding two more boxes over the next 10 sessions, a few more individuals acquired the task in each group, suggesting that acquiring the task was also related to the number of available boxes. Thus, the timing of acquisition was clearly influenced by this design, violating the assumption of the *TADA*.

### Model implementations

All analyses were conducted using R (v. 4.4.2; R Core team, 2024). To test the significance of fixed effect predictors as a whole, we compared the fit of the full model with that of the null model comprising only random or control factors (Schielzeth and Forstmeier 2009; Forstmeier and Schielzeth 2010). We obtained confidence intervals for all models by means of parametric bootstraps using the function ‘bootMer’ of the package ‘lme4’ (Bates et al. 2015), applying 1000 bootstraps. We checked for collinearity issues by determining variance inflation factors for a standard linear model without random effects using the package ‘car’ (Fox and Weisberg 2019). To estimate model stability, we dropped levels of the random effects one at a time from the dataset and compared the obtained estimates to the estimates obtained for the full dataset.

### Ethical note

This study adhered to the ASAB/ABS Guidelines for the Treatment of Animals in Behavioural Research and Teaching and to the legal requirements of the country (Madagascar) in which the study was carried out. The protocol for this research was approved by the Malagasy Ministry of the Environment, Water, and Forests (064,22/ MEDD/ SG/ DGGE/ DAPRNE/ SCBE.Re). Research on ring-tailed lemurs was authorised by the family de Heaulme, the owners of Berenty Private Reserve. Although our experiments involved some feeding competition, we tried to minimize it by conducting the experiments only for four successive days. In addition, we did not observe any changes in the behaviour of the groups after conducting the experiments.

## Results

### Number of learners and speed of learning

In red-fronted lemurs, the proportion of individuals that learned to open the food box irrespective of the technique used, 47±21% (mean ± SD), did not differ from the proportion of individuals that did not learn the task (proportion test: χ^2^=0.281, df=1, p=0.596). In ring-tailed lemurs, however, the proportion of individuals learning the task was with 28±4% (mean ± SD) lower in comparison to the proportion of individuals that did not learn the task (proportion test: χ^2^=4.367, df=1, p=0.037). The probability of learning to open the box irrespective of the technique used was not influenced by sex or age class and only by trend by the interaction between species and condition, i.e., groups with or without demonstrators (Table 1; likelihood ratio test full-null model: χ^2^=10.33, df=4, p=0.035).

**Table 1.** Factors influencing the probability of learning to open the box irrespective of the technique used in red-fronted and ring-tailed lemurs.

| Term | Estimate | SE | P-value |
| --- | --- | --- | --- |
| Intercept | -0.961 | 0.714 | <sup>1</sup> |
| Demonstrator (yes) | -1.625 | 0.790 | 0.040 |
| Species (ring-tailed lemurs) | -2.321 | 0.942 | 0.014 |
| Sex (female) | 0.936 | 0.552 | 0.090 |
| Age class (juvenile) | 0.446 | 0.799 | 0.576 |
| Species (ring-tailed lemurs): demonstrator (yes) | 2.204 | 1.159 | 0.057 |

<sup>1</sup> Not shown as has no meaningful interpretation

Red-fronted lemurs learned the task on average in 1.07 ± 0.27 (mean±SD) trials and thus more quickly than ring-tailed lemurs, which needed 2.60 ± 2.06 trials to open the food box successfully (Figure 2; Mann-Whitney *U* test: *U* = 31.5, *N*_1_ = 14, *N* _2_ = 10, *P* = 0.005). The presence of a demonstrator did not influence the learning speed in both species (Mann-Whitney *U* test: red-fronted lemurs: *U* = 21, *N*_1_ = *N* _2_ = 7, *P* = 0.391; ring-tailed lemurs: *U* = 7.5, *N*_1_ = 7, *N* _2_ = 3, *P* = 0.551).

**Figure 2.**
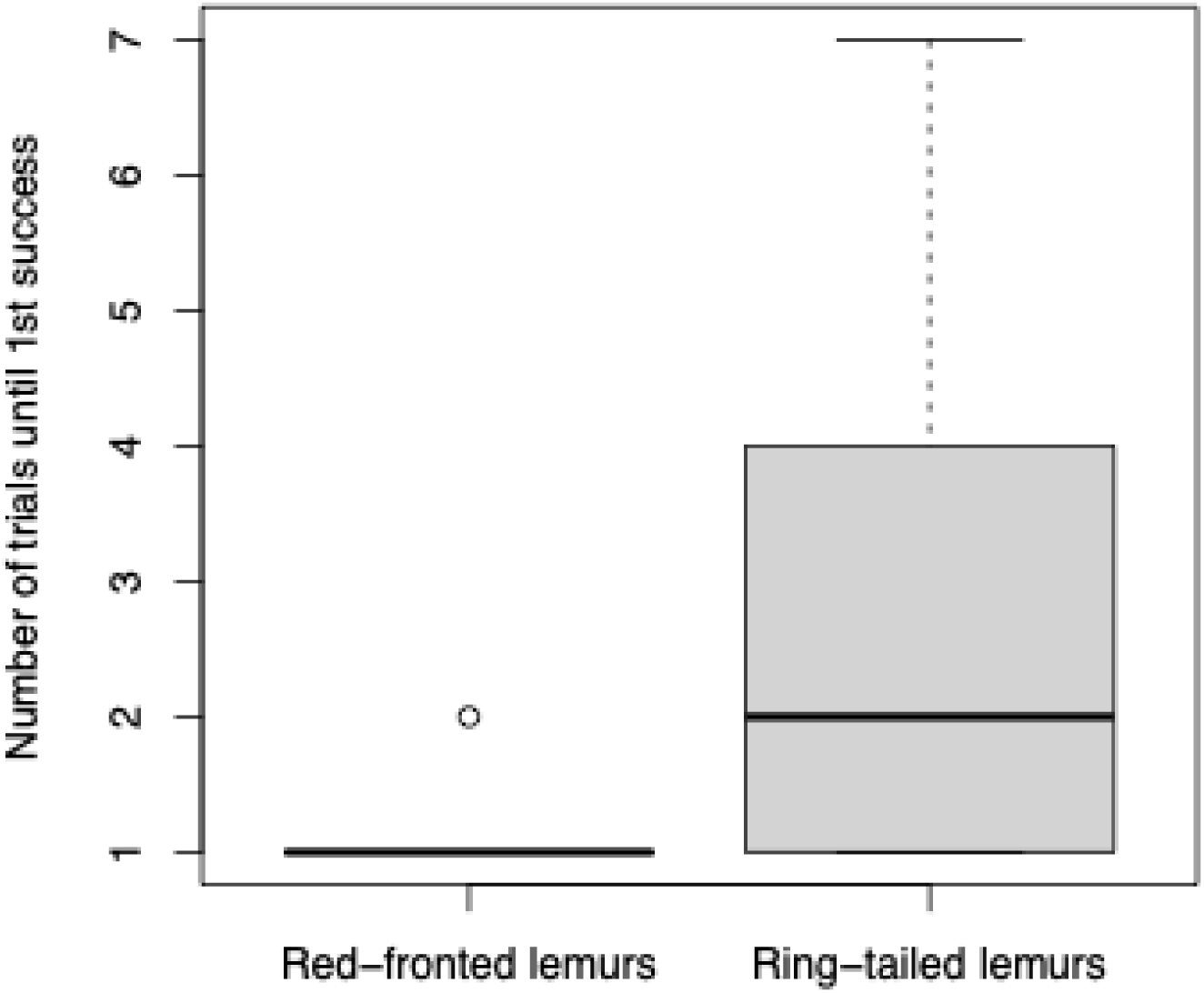
Number of trials an individual needed to open the food box for the first time. Red-fronted lemurs (EF) required significantly less trials to learn to open the food box compared to ring-tailed lemurs (LC).

### Homogeneity in the spread of techniques

All learners discovered both techniques to open the boxes. The proportion of using the lifting technique differed between treatments in both species (full-null model: red-fronted lemurs: F=519.88_2_, *P*<0.001; ring-tailed lemurs: F=20.14_2_, *P*<0.001; Table 3a, b). In red-fronted lemurs, individuals in groups in which one of the techniques was seeded performed more often the seeded technique and individuals in open groups performed more often the lifting technique (Figure 3). However, ring-tailed lemurs performed overall more often the lifting technique. Individuals in the group in which the pushing technique was seeded and in the open group performed more often the lifting technique than individuals in the group in which the lifting technique was seeded (Figure 3; Table 3b).

**Figure 3.**
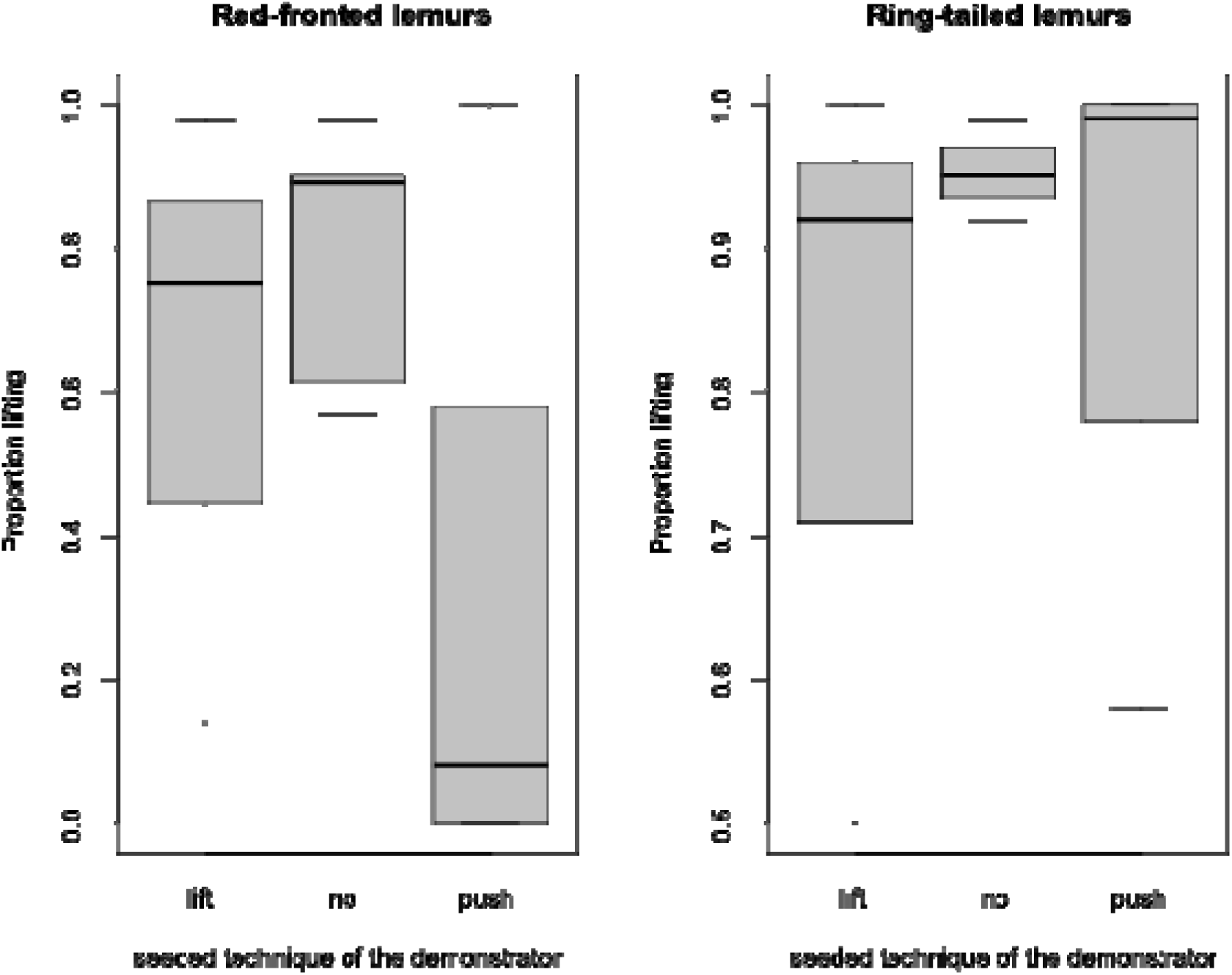
Proportion in the usage of the lifting technique for groups in which the lifting-, no-, or pushing technique was seeded in red-fronted and ring-tailed lemurs.

**Table 3.** Factors influencing the proportion of using the lifting technique in a) red-fronted and b) ring-tailed lemurs as well as factors influencing the proportion of scrounging events in c) red-fronted and d) ring-tailed lemurs.

| Species | Term | Estimate | SE | P-value |
| --- | --- | --- | --- | --- |
| a) Red-fronted lemurs | Intercept | 3.178 | 0.255 | <sup>1</sup> |
|  | Demonstrator (no) | -1.414 | 0.266 | <0.001 |
|  | Demonstrator (push) | -4.839 | 0.316 | <0.001 |
| b) Ring-tailed lemurs | Intercept | 1.981 | 0.533 | <sup>1</sup> |
|  | Demonstrator (no) | 1.276 | 0.548 | 0.020 |
|  | Demonstrator (push) | 2.472 | 0.630 | <0.001 |
| c) Red-fronted lemurs | Intercept | -2.238 | 0.770 | <sup>1</sup> |
|  | Technique (push) | 0.517 | 0.161 | 0.001 |
|  | Session | 0.278 | 0.178 | 0.120 |
|  | Sex (male) | 0.109 | 0.563 | 0.846 |
|  | Age class (juvenile) | -0.221 | 0.876 | 0.801 |
| d) Ring-tailed lemurs | Intercept | -4.263 | 0.440 | <sup>1</sup> |
|  | Technique (push) | 0.237 | 0.461 | 0.607 |
|  | Session | 0.662 | 0.221 | 0.003 |
|  | Sex (male) | -0.722 | 0.601 | 0.229 |
|  | Age class (juvenile) | 1.264 | 0.652 | 0.052 |

<sup>1</sup> Not shown as has no meaningful interpretation

These results may suggest that the lifting technique might have been easier to perform than the pushing technique. However, in both species, the proportion of unsuccessful actions was higher when using the lifting than the pushing technique (red-fronted lemurs: lifting technique: 0.25 ± 0.52 % (mean ±SD), pushing technique: 0.04 ± 0.05 %, Wilcoxon signed-ranks test: T=4, *N*=14, *P*=0.019; ring-tailed lemurs: lifting technique: 0.54 ± 0.91 % (mean ±SD), pushing technique: 0.75 ± 0.15 %, T=3.5, *N*=10, *P*=0.049), suggesting that the lifting-technique might have been indeed more difficult.

In both species, individuals also scrounged. In red-fronted lemurs in 17.41% of successful events, group members scrounged, which was about eight times higher than in ring-tailed lemurs where only in 2.17% of successful events a group member scrounged (proportion test: χ^2^=499.46_1_, *P*<0.001). In red-fronted lemurs, the proportion of scrounging events was higher when individuals used the pushing technique (full-null model: red-fronted lemurs: χ^2^=11.58, *P*=0.003; Figure 4, Table 3c), though this technique seemed to be more difficult. Sex and age class of the producer did not influence the proportion of scrounging events (Table 3c). In ring-tailed lemurs, however, the used technique, age and sex of the producer did not influence the proportion of scrounging events (full-null model: red-fronted lemurs: χ^2^=3.77, *P*=0.151; Table 3d). Session covaried positively with the proportion of scrounging events increasing over time (Table 3d). Since the full-null model comparison was not significant, these results should be considered carefully.

**Figure 4:**
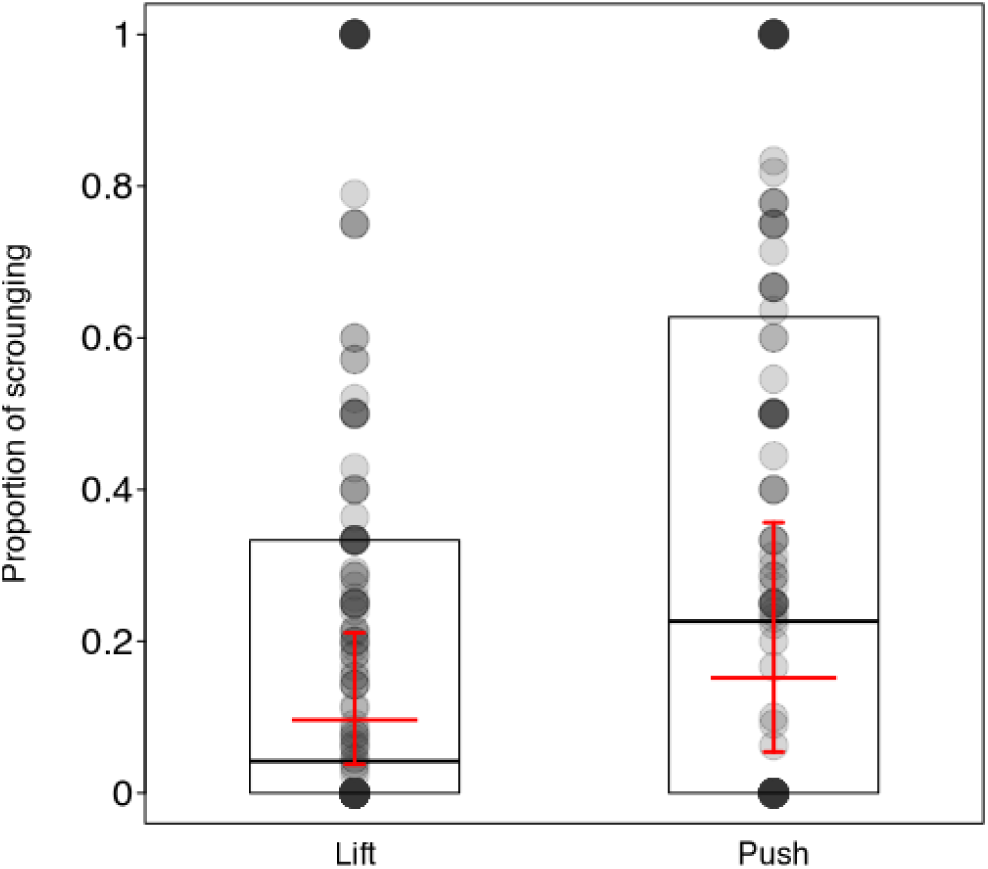
The proportion of scrounging events for each technique used in red-fronted lemurs. Boxes present boxplots depicting the 1. and 3. Quartile. Black lines present the median. Red horizontal lines present the estimates and vertical lines the 95% confidence intervals.

### Order of acquisition diffusion analysis

In red-fronted lemurs, two *OADA* models provided the best fits with both revealing significant support for social transmission (Table 4). In the first *OADA* model, social transmission was kept constant across groups and individual-level variables were not included (AICc=47.29, p=0.011). In the second *OADA* model, the magnitude of social transmission was set constant within conditions, i.e. demonstrator and open groups, but allowed to vary between conditions (AICc=47.51, p=0.004). The models in which we added individual level variables, aggression rates and sex, did not provide a better fit (Table 4). In ring-tailed lemurs the *OADA* model with social transmission kept constant, excluding individual-level variables provided the best fit, revealing significant support for social learning (AICc=45.80, p=0.042, Table 4). In contrast to red-fronted lemurs the model in which the magnitude of social transmission was set constant within conditions but allowed to vary between conditions, i.e. demonstrator and the open group, did not reveal a better fit (AICc=56.81). As in red-fronted lemurs, the models in which we added individual level variables did not provide a better fit (Table 4).

**Table 4.** Results of the 6 OADA models of red-fronted and ring-tailed lemurs, respectively. In bold: models with the best fit (lowest AICc). ST= Transmission, ILVs: Individual level variables (sex, aggression rates)

| **OADA: red-fronted lemurs** |  | **Df** | **LogLik** | **AIC** | **AICc** | **LR** | **p** | **ST in the 4 groups** |
| --- | --- | --- | --- | --- | --- | --- | --- | --- |
| 1. ST constant across the 4 groups | with ST | 1 | 22.479 | 46.958 | **47.291** | 6.460 | 0.011 | 1,1,1,1 |
|  | without ST | 0 | 25.709 | 51.418 | 51.418 |  |  |  |
| 2. ST constant across the 4 groups, including ILVs | with ST | 3 | 20.838 | 47.676 | 50.076 | 4.602 | 0.032 | 1,1,1,1 |
|  | without ST | 2 | 23.139 | 50.279 | 51.370 |  |  |  |
| 3. ST varies between the 4 groups | with ST | 4 | 21.451 | 50.902 | 52.235 | 8.516 | 0.074 | 1,2,3,4 |
|  | without ST | 0 | 25.709 | 51.418 | 51.418 |  |  |  |
| 4. ST varies between the 4 groups, including ILVs | with ST | 6 | 16.435 | 44.870 | 56.87 | 13.409 | 0.009 | 1,2,3,4 |
|  | without ST | 2 | 23.139 | 50.279 | 51.37 |  |  |  |
| 5. ST constant within conditions (demonstrator, open groups) but varying between conditions | with ST | 2 | 21.457 | 46.915 | **47.582** | 8.503 | 0.014 | 1,1,2,2 |
|  | without ST | 0 | 25.709 | 51.418 | 51.418 |  |  |  |
| 6. ST constant within conditions (demonstrator, open groups) but varying between conditions | with ST | 4 | 20.050 | 48.100 | 52.545 | 6.179 | 0.045 | 1,1,2,2 |
|  | without ST | 2 | 23.139 | 50.279 | 51.37 |  |  |  |
| **OADA: ring-tailed lemurs** |  |  |  |  |  |  |  | **ST in the 3 groups** |
| 1. ST constant across the 4 groups | with ST | 1 | 20.681 | 43.362 | **43.862** | 4.121 | 0.0423 | 1,1,1 |
|  | without ST | 0 | 22.742 | 45.484 | 45.484 |  |  |  |
| 2. ST constant across the 4 groups, including ILVs | with ST | 3 | 20.320 | 46.640 | 50.640 | 2.664 | 0.103 | 1,1,1 |
|  | without ST | 2 | 21.652 | 47.304 | 49.018 |  |  |  |
| 3. ST varies between the 4 groups | with ST | 3 | 19.151 | 44.303 | 45.803 | 7.1809 | 0.066 | 1,2,3 |
|  | without ST | 0 | 22.742 | 45.484 | 45.484 |  |  |  |
| 4. ST varies between the 4 groups, including ILVs | with ST | 5 | 18.907 | 47.815 | 62.815 | 5.4894 | 0.139 | 1,2,3 |
|  | without ST | 2 | 21.652 | 47.304 | 49.018 |  |  |  |
| 5. ST constant within conditions (demonstrator,<br>open groups) but varying between conditions | with ST | 2 | 20.563 | 45.126 | 46.126 | 4.358 | 0.113 | 1,2,1 |
|  | without ST | 0 | 22.742 | 45.484 | 45.484 |  |  |  |
| 6. ST constant within conditions (demonstrator,<br>open groups) but varying between conditions | with ST | 4 | 20.404 | 48.808 | 56.808 | 2.4963 | 0.287 | 1,2,1 |
|  | without ST | 2 | 21.652 | 47.304 | 49.018 |  |  |  |

### Network efficiency

Across species, social network metrics influenced the proportion of learners in these lemurs. Social tolerance (Figure 4) and global efficiency correlating positively with the proportion of learners (Spearman rank correlation: social tolerance: r_S_=0.85, N=7, P=0.008; global efficiency: r_S_ =0.9, N=6, P=0.007), whereas average efficiency did not correlate with the proportion of learners (Spearman rank correlation: r_S_=-0.46, N=6, P=0.823).

**Figure 4.**
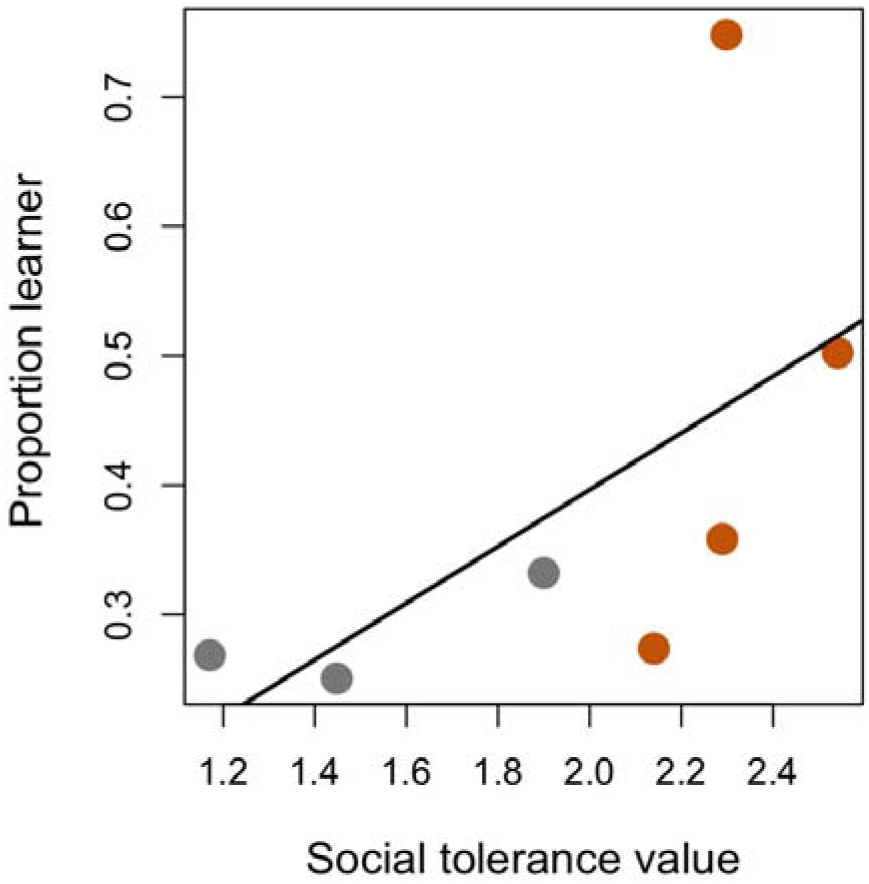
Proportion of learners per group as a function of social tolerance taken from co-feeding experiments. Though ring-tailed lemurs (grey dots) were less tolerant than red-fronted lemurs (orange dots), the correlation between social tolerance and proportion of learners within a group was observable across both species.

## Discussion

We presented two-option feeding boxes in a standardized set-up, either with or without a trained demonstrator, to examine social learning of a novel behaviour in seven groups of two wild lemur species (n=4 red-fronted lemurs; n=3 ring-tailed lemurs). We also measured social tolerance using an established co-feeding paradigm. We investigated whether the new behaviours would spread further, more homogenously, and more quickly in the more tolerant species compared to the less tolerant species. Overall, red-fronted lemurs were more likely to learn how to open the food boxes and learned it faster compared to ring-tailed lemurs. In both species, we found support for social learning, which was independent of whether there was a seeded demonstrator or not. All individuals used both techniques to open the food boxes, but had a preference for one of the techniques. The presence of a demonstrator of the non-preferred technique influenced the frequency of the used technique in the red-fronted-but not the ring-tailed lemurs. In both species, individuals scrounged from other successful individuals. In the more tolerant red-fronted lemurs, the proportion of scrounging events was higher than in the less tolerant ring-tailed lemurs. Red-fronted lemurs scrounged proportionally more often when the pushing technique was used. Lastly, social tolerance and global efficiency facilitated learning in both species.

### Number of learners and learning speed

In red-fronted lemurs, the proportion of individuals who solved the problem or not did not significantly differ from each other. In ring-tailed lemurs, however, the proportion of solvers was significantly lower than the number of non-solvers. Our results fit with previous findings showing that information spreads more homogeneously within red-fronted lemurs and only to subgroups in ring-tailed lemurs (Kendal et al., 2010; Schnoell & Fichtel, 2012). Accordingly, on average, red-fronted lemurs learned the solution faster than ring-tailed lemurs. Though it might be possible that the problem was more difficult for ring-tailed lemurs than it was for red-fronted lemurs, we doubt this explanation. Previous studies have shown that ring-tailed lemurs are capable problem solvers and are qualitatively equal in the physical cognitive domain compared to other lemurs (Fichtel et al., 2020; Kittler et al., 2015). Hence, it is more likely that within the groups of ring-tailed lemurs, fewer individuals had access to the food boxes, which on average increased the latency to solve and ultimately decreased the probability of solving the problem at all. This explanation is coherent with the idea that ring-tailed lemurs are less socially tolerant than red-fronted lemurs and show higher aggression rates when resources can be monopolized (Fichtel et al., 2018). In addition, in red-fronted lemurs, more individuals during more events were able to scrounge than in ring-tailed lemurs, supporting the notion that scrounging is more prevalent in the more socially tolerant species.

We did not find an effect of age or sex on the likelihood of learning, supporting similar findings in red-fronted lemurs (Schnoell & Fichtel 2012; Karakoc et al., 2025). Age appears to have different effects on innovation, with in some species, younger individuals tending to be more explorative and develop novel foraging techniques (Kummer & Goodall, 1997; Nishida, et al., 1983; Perry, 2020; Perry et al., 2017), the trend seems to be reversed in other species like for instance, in marmosets (Kendal et al., 2005; but see Sehner et al., 2022). Then again, in meerkats, younger individuals tend to be more explorative during foraging tasks, but they often fail to innovate a solution, and the majority of the innovations are performed by adult individuals (Thornton & Samson, 2012). Surprisingly, sex did not significantly influence the probability of innovating. Though this result fits the egalitarian system of and earlier findings in red-fronted lemurs (Karakoc et al., 2025), one would expect a significant difference within the more despotic and female-dominated ring-tailed lemurs (Kappeler et al., 2022), which was also not the case. A likely explanation for this is may the younger male ring-tailed lemurs who were among the solving individuals, were still tolerated by their mothers and thus gained access to the food boxes. Thus, the effect of sex might have been masked by these younger ring-tailed lemur males.

### Homogeneity in the spread of techniques

In both species, all individuals who learned to open the food box were capable of performing both techniques in the end. The presence of a demonstrator affected the two species differently. Though only as a trend, the presence of a demonstrator had a stronger effect on ring-tailed lemurs than on red-fronted lemurs. Nevertheless, both species seem to socially learn the solution, either from the seeded demonstrator or from independent first innovators (see below). The main difference between the two species was the frequency with which they used one or the other technique. Whereas ring-tailed lemurs showed a clear tendency to use the lifting solution, irrespective of the seeded technique, red-fronted lemurs showed a higher tendency toward the pushing technique if that was the seeded technique. It appears that ring-tailed lemurs socially learn via simple mechanisms of social enhancement (O’Mara & Hickey, 2012) which draw their attention toward an object or location but not to the actions of a demonstrator.

Alternatively, since ring-tailed lemurs also learned the pushing technique, it might be that they find this technique simpler and thus preferable. Though lemurs generally lack a precision grip and have inferior dexterity compared to haplorrhines (Torigoe 1985; Kittler et al., 2015), neither technique required a precision grip. The fact that individuals made more mistakes using this technique does not support the idea that lifting was easier and thus raises the question of why they preferred this technique in all conditions.

Red-fronted lemurs, like ring-tailed lemurs, used mainly the lifting technique, independent of whether the lifting technique was seeded or not. However, red-fronted lemurs performed the pushing technique more often than the lifting technique if pushing was seeded by a demonstrator. Again, all solvers were capable of using both techniques, and lifting produced more mistakes, raising the question of why red-fronted lemurs behaved differently than ring-tailed lemurs? The proportion of scrounging events was higher when red-fronted lemurs used the pushing technique. Therefore, red-fronted lemurs may use the lifting technique more often to avoid scroungers, even though it appears to be more difficult. Interestingly, a recent study of the same population found that the number of scroungers was independent of the technique used by the individual opening a similar feeding box as in this study (Karakoç et al., 2025). This suggests that the preference for lifting may be driven by scrounging itself, rather than by the number of individuals scrounging simultaneously. Although the pushing technique facilitates scrounging, individuals in groups where pushing was introduced exhibited a preference for it. Therefore, an alternative, non-mutually exclusive explanation could be that red-fronted lemurs in this group exhibit greater conformity in their foraging behaviour. Such normative conformity, i.e. conformity motivated by the desire to maintain group cohesion, has previously been demonstrated in vervet monkeys (van de Waal et al., 2013). Although formal tests are required to demonstrate conformity in red-fronted lemurs, our results suggest that a concept traditionally associated with humans (Whiten et al., 2011) and hypothesised to be crucial for cultural evolution may have existed since the beginning of the primate lineage (see also Aplin, 2019, regarding birds).

In contrast, in the less tolerant ring-tailed lemurs, scrounging occurred less often, and the technique used did not influence the proportion of scrounging events. Hence, scrounging does not explain the preference for the lifting technique. Alternatively, also in ring-tailed lemurs, conformity in foraging behaviour may explain this preference.

### Order of acquisition and network efficiency

We found support for social learning in both species and the best model for both species considered social learning equal across conditions. Hence, it is not the presence of a seeded demonstrator that affects social learning, but rather whether any individual can serve as a role model. Our results fit with previous findings that, in both ring-tailed and red-fronted lemurs, information transfer occurs socially rather than through independent learning (Kittler et al., 2015; O’Mara & Hickey, 2012; Schnoell et al., 2014; Schnoell & Fichtel, 2012, but see Huebner & Fichtel, 2015). Furthermore, our results fit in the general perception that the social network is a robust predictor for the spread of information (Allen et al., 2013; Claidière et al., 2013; Hobaiter et al., 2014; Wild et al., 2019). Consistent with the results on initial problem-solving acquisition, individual traits like sex and aggression had no significant effect on the likelihood of information being transferred socially.

Social tolerance and global network efficiency, but not average network efficiency, predicted the proportion of learners per group. This pattern suggests that tolerance maintains cohesion on a group level, allowing for at least weak connections between individuals and thus information transfer. Our results confirm the findings of Pasquaretta and colleagues (2014) that on a global level, more egalitarian societies are more efficient than more hierarchical ones. It also supports the notion that social tolerance, even if measured in a different context than the one relevant for learning, can predict social information transfer (Coelho et al., 2024; Garcia-Nisa et al., 2023; Whiten & van Schaik, 2007). Although tolerance levels can vary within species depending on factors such as resource monopolizability (White et al., 2007), ecological conditions (de Oliveira Terceiro et al., 2021; Fichtel et al., 2018), and group size (Berghänel et al., 2025) it nevertheless remains a reliable proxy for general information transfer (Coelho et al. 2024).

In conclusion, tolerance plays a vital role in shaping the dynamics of social learning in wild red-fronted and ring-tailed lemurs. The more egalitarian red-fronted lemurs demonstrated to learn the solution to a problem faster, and involved a greater proportion of group members compared to the more despotic ring-tailed lemurs. While both species showed evidence of social learning, irrespective of the presence of a seeded demonstrator, the type of technique adopted was influenced by species-specific social structures. These findings provide important insights into how social dynamics influence social learning, highlighting the importance of group cohesion in facilitating information flow. Overall, our findings support the view that social tolerance promotes social learning and can serve as a critical driver of group-specific behaviours, hence contributing to a broader understanding of

## Acknowledgment

We thank A.V. Schnöll for collecting some of the data and W. Hoppitt for support in the OADA-analyses. This publication greatly benefited from discussions of the Collaborative Research Center “Cognition of Interaction” funded by the Deutsche Forschungsgemeinschaft (DFG, German Research Foundation) – Project-ID 454648639 – SFB 1528. The authors thank the Swiss National Foundation (SNF) for its support. S. Sehner was supported by a PostDoc Mobility Grant (grant number P500PB_217864/1).

## Author contributions

SS: writing – original draft; FR: editing original draft; CF: conceptualization, data curation, formal analysis, investigation, methodology, project administration, validation, visualization, writing – original draft.

## Data availability

Date and R code are available under https://figshare.com/s/6237c92293500c2df636

